# A Drug Combination Approach Targeting Both Growing Bacteria and Dormant Persisters Eradicate Persistent *Staphylococcus aureus* Biofilm Infection

**DOI:** 10.1101/686097

**Authors:** Rebecca Yee, Yuting Yuan, Andreina Tarff, Cory Brayton, Naina Gour, Jie Feng, Wanliang Shi, Ying Zhang

## Abstract

*Staphylococcus aureus* can cause a variety of infections, many of which involve biofilm infections. Inside biofilms, growing and non-growing bacteria such as persisters co-exist, making it challenging to completely eradicate a persistent and recurrent infection with current treatments. Despite the clinical relevance, most of the current antibiotic treatments mainly kill the growing bacteria and have poor activity against non-growing persister bacteria and thus have limited effect on treating persistent infections including biofilm infections. We previously proposed a Yin-Yang model using a drug combination approach targeting both growing bacteria and persister bacteria for more effective clearance of persistent infections. Here, as a proof of principle, we showed that combining drugs that have high activity against growing forms, such as vancomycin or meropenem, with drugs that have robust anti-persister activity, such as clinafloxacin and oritavancin, could completely eradicate *S. aureus* biofilm bacteria in vitro. In contrast, single or two drugs including the current treatment for persistent *S. aureus* infection doxycycline plus rifampin failed to kill all biofilm bacteria in vitro. We then developed a chronic persistent skin infection mouse model with biofilm-seeded bacterial inocula demonstrating that biofilm bacteria caused more severe and persistent skin lesions than log phase *S. aureus* bacteria. More importantly, we found that the drug combination which eradicated biofilm bacteria in vitro is more efficacious than current treatments and completely eradicated *S. aureus* biofilm infection in mice. The complete eradication of biofilm bacteria is attributed to the unique high anti-persister activity of clinafloxacin, which could not be replaced by other fluoroquinolones such as moxifloxacin, levofloxacin or ciprofloxacin. Our study is the first to demonstrate that the combination of meropenem, daptomycin, plus clinafloxacin completely cleared the persistent infection, healed the lesions, and had less inflammation, while mice treated with doxycycline plus rifampin, the current clinically recommended treatment for chronic tissue infection, failed to do so. We also compared our persister drug combination with other approaches for treating persistent infections including gentamicin+fructose and ADEP4+rifampin in the *S. aureus* biofilm infection mouse model. Neither gentamicin+fructose nor ADEP4+rifampin could eradicate or cure the persistent biofilm infection in mice. In contrast, our drug combination regimen with persister drug clinafloxacin plus meropenem and daptomycin completely eradicated and cured the persistent biofilm infection in 7 days. An unexpected observation is that ADEP4 treatment group developed worsened skin lesions and caused more extensive pathology than the untreated control mice. Our study demonstrates an important treatment principle for persistent infections by targeting both growing and non-growing heterogeneous bacterial populations utilizing persister drugs for more effective eradication of persistent and biofilm infections. Our findings may have implications for improved treatment of many other persistent infections in general.

## Introduction

Methicillin-resistant *Staphylococcus aureus* (MRSA) strains are highly prevalent in healthcare and community-acquired staphylococcal infections ^1^. The mortality rate associated with MRSA infection is as high as 40% ^2,3^. As an opportunistic pathogen, *S. aureus* is the most common cause of skin infections and can also cause chronic infections such as endocarditis, osteomyelitis, and prosthetic joint infections ^4-6^. In particular, indwelling devices are conducive to biofilm formation, complicating treatment and leading to prolonged infections. Globally, persistent and chronic infections are a huge burden to public health as they increase the length of hospital stay, cause relapse, cost of treatment, and risk of death by at least three-folds^7^.

Bacteria in biofilms are more tolerant to antibiotics compared to planktonic cells^8^. Studies have shown that antibiotics do indeed penetrate the biofilm but they do not always kill the bacteria, suggesting that tolerance to treatment is not due to impaired antibiotic penetration or genetic resistance ^9,10^, but due to dormant non-growing or slowing growing persister bacteria. Bacteria inside the biofilm are quite heterogeneous as some cells grow slowly, which are representative of stationary phase bacteria, while others form dormant persister cells due to the high cell density, nutrient and oxygen limiting environment inside the biofilm matrix ^11^.

First described in 1942, Hobby et al. found that while 99% of *S. aureus* cells were killed by penicillin about 1% of residual metabolically quiescent or dormant cells called persister cells were not killed ^12^. The persister cells were not resistant to penicillin and hence, did not undergo genetic changes but were phenotypic variants that became tolerant to antibiotics ^13^. Similarly, a clinical observation was also made as penicillin failed to clear chronic infections due to the presence of persister cells found in patients ^13^. While the mechanisms of *S. aureus* persistence were largely unknown for a long time, recent studies have shown that pathways involved in quorum sensing, pigmentation production, and metabolic processes such as oxidative phosphorylation, glycolysis, amino acid and energy metabolism ^14-18^ are responsible. Understanding the pathways of persistence would facilitate development of novel drugs and therapeutic approaches to more effectively eradicate persistent bacterial infections.

Despite the observation of persister bacteria from 1940s and their implications in causing prolonged treatment and post-treatment relapse, the importance of persister bacteria in clinical setting has been ignored largely because no persister drugs have been found that can shorten treatment duration and reduce relapse in clinically relevant persistent infections. The importance of persister drug to more effectively cure persistent infections is only recognized recently in the case of tuberculosis persister drug pyrazinamide (PZA) which shortens the tuberculosis treatment from 9-12 months to 6 months after its inclusion in a drug combination setting^19^. PZA’s activity in killing persisters, unlike the other drugs used to treat tuberculosis, is crucial in developing a shorter treatment ^19-22^. A drug like PZA validates an important principle of use of a persister drug in combination with other drugs targeting both persisters and growing cells in formulating an effective therapy for chronic persistent infections ^23-25^. More recently, a similar approach has been used to identify effective drug combinations to eradicate biofilm-like structures consisting of heterogeneous cells of *Borrelia burgdorferi in vitro* ^26^.

Using this approach, in a recent study aimed at identifying drugs targeting non-growing persisters, we used stationary phase culture of *S. aureus* as a drug screen model and identified several drugs such as clinafloxacin and tosufloxacin with high activity against *S. aureus* persisters ^27^. However, their activities alone and in drug combinations in killing biofilms have not been evaluated in vitro or in related infections caused by *S. aureus* in vivo. In this study, we developed drug combinations that can more effectively eradicate *S. aureus* biofilms by formulating drug combinations that have high activities against growing bacteria and non-growing persisters in a biofilm model in vitro initially. Then, we established a persistent skin infection mouse model for *S. aureus* using “biofilm seeding” and evaluated drug combinations in clearing the biofilm infection in this persistent skin infection model. Here, we show that combining meropenem and daptomycin targeting growing bacteria, with clinafloxacin targeting persister bacteria led to complete eradication of *S. aureus* biofilm not only *in vitro* but more importantly also *in vivo* in a murine model of persistent skin infection, whereas other approaches for treating persistent infections such as aminoglycoside plus sugar or ADEP4 plus rifampin and the currently recommended drug combination treatment without persister drugs failed to do so.

## Results

### Commonly used treatments for MRSA have poor activity against biofilms in vitro

While vancomycin is highly effective in killing MRSA bacteria in vitro, monotherapy with vancomycin may not be the most effective in clearing chronic infections with *S. aureus*. For conditions such as osteomyelitis and prosthetic joint infections, treatment with vancomycin as a monotherapy or drug combination for at least 6 weeks are recommended. Drug combination such as doxycycline + rifampin for up to 10 days, vancomycin + gentamicin + rifampin for at least 6 weeks are recommended to treat chronic infections such as recurrent tissue infections and endocarditis on prosthetic valves, respectively (The Johns Hopkins Antibiotics Guide). We first evaluated the activity of the above drugs in killing biofilm bacteria *in vitro* using traditional bacterial cell counts (Fig. 1A), viability assessment by SYBR Green I/Propidium Iodide staining that has been developed to screen for drugs targeting borrelia persister bacteria ^28^ (Fig. 1B), and staining of absolute biofilm (Fig. 1C-G). We found that such clinically used combinations are not completely effective against biofilms. After 4-day treatment, biofilm bacteria were not completely eradicated by any of current treatments with vancomycin alone, or doxycycline + rifampin or vancomycin+gentamicin+rifampin as shown by significant numbers of bacteria remaining (Fig. 1). However, it is worth noting that vancomycin+gentamicin+rifampin was more active than vancomycin alone or doxycycline + rifampin in killing biofilm bacteria (Fig. 1A).

**Figure 1.**
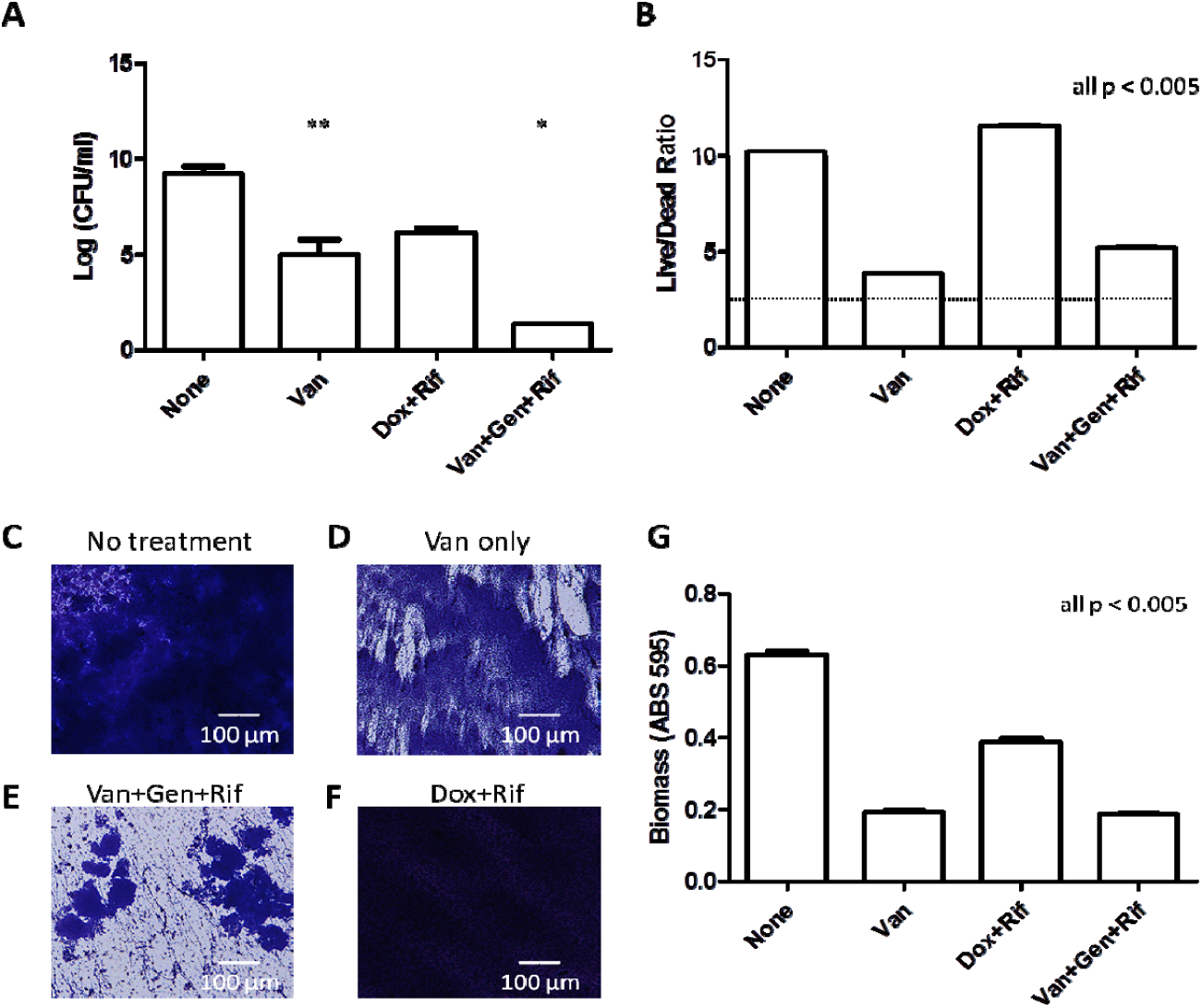
Clinically recommended treatments for chronic *S. aureus* infections only partially killed biofilm bacteria *in vitro*. Treatments (all 50 μM) of vancomycin alone, doxycycline+rifampin, and the combination of vancomycin+gentamicin+rifampin for 4 days were evaluated for biofilm killing by CFU enumeration (A) and viability staining using SYBR Green I/PI (B). Images of biofilm biomass (C-F) and quantification of absolute biofilm biomass (G) after respective antibiotic treatments. Vancomycin, Van; Doxycycline, Dox; Rifampin, Rif; Gentamicin, Gen. Student’s *t-*test, *p□<□0.05, **p□<□0.005, ***p□<□0.0005.

### Identification of drug combinations with strong anti-biofilm activity

To address the clinical unmet need of better treatments against persistent infections, we hypothesize that a drug combination that includes drugs that act on growing bacteria such as cell wall (e.g. vancomycin, meropenem) or cell membrane inhibitors (e.g. daptomycin) plus a drug that acts on persister bacteria will be a more potent drug combination in eradicating biofilm bacteria. Previous studies from our lab identified tosufloxacin and clinafloxacin as having strong anti-persister activity against *S. aureus* ^27^. In order to identify a potent combination, we tested various drug combinations that include drugs against both growing bacteria and non-growing persisters in an *in vitro* biofilm model. Biofilms of *S. aureus* strain USA300, a common circulating strain of community acquired-MRSA (CA-MRSA), were grown in 96-well microtiter plates to allow biofilm formation on the bottom of the wells^29^. While we previously showed that tosufloxacin had robust activity against *S. aureus* persister cells, drug combination of vancomycin/meropenem + daptomycin + tosufloxacin achieved only partial eradication, with 10^5^ CFU/ml in biofilms remaining after treatment (Fig. 2A-B). In contrast, combination of vancomycin/meropenem + daptomycin + clinafloxacin showed absolute eradication of biofilms after 4-day treatment as shown by 0 CFU and a live/dead ratio below the limit of detection (Fig. 2A-B). Although we used the same molar concentration of each drug (50 μM of each drug) in our drug screen for comparison of relative drug activity, to evaluate the activity of the combination in a more clinically relevant manner, we treated the biofilms with the drugs at their Cmax concentrations (Table 1). Our findings with Cmax drug concentrations were confirmatory as the combination of vancomycin/meropenem + daptomycin + clinafloxacin still achieved complete eradication while our no treatment control and the clinically used combination of doxycycline + rifampin could not. The clearance of biofilms was confirmed by both CFU counts and viability staining (Fig. 2C-D).

**Table 1.** Drug Dosage, Scheduling, and Administration.

| **Drug** | **Dosage** | **Route** | **Times Treated** | **Cmax tested (clinically achievable concentrations)** |
| --- | --- | --- | --- | --- |
| **Vancomycin** | 110 mg/kg | Intraperitoneally | Twice/daily | 20 µg/ml |
| **Daptomycin** | 50 mg/kg | Intraperitoneally | Once/daily | 80 µg/ml |
| **Meropenem** | 50 mg/kg | Intraperitoneally | Once/daily | 20 µg/ml |
| **Clinafloxacin** | 50 mg/kg | Intraperitoneally | Once/daily | 2 µg/ml |
| **Doxycycline** | 100 mg/kg | Oral | Twice/daily | 5 µg/ml |
| **Rifampin** | 10 mg/kg | Oral | Twice/daily | 5 µg/ml |
| **Moxifloxacin** | 100 mg/kg | Oral | Once/daily | 4 µg/ml |
| **ADEP4** | 25 mg/kg and 35 mg/kg | Intraperitoneally | Twice/daily |  |
| **Rifampin (for ADEP4 combination)** | 30 mg/kg | Intraperitoneally | Once/daily |  |
| **Gentamicin** | 20 mg/kg | Intraperitoneally | Once/daily |  |
| **Fructose** | 1.5 g/kg | Intraperitoneally | Once/daily |  |
| **Oritavancin** | --- | --- | --- | 5 µg/ml |
| **Ciprofloxacin** | --- | --- | --- | 10 µg/ml |
| **Levofloxacin** | --- | --- | --- | 10 µg/ml |

**Figure 2.**
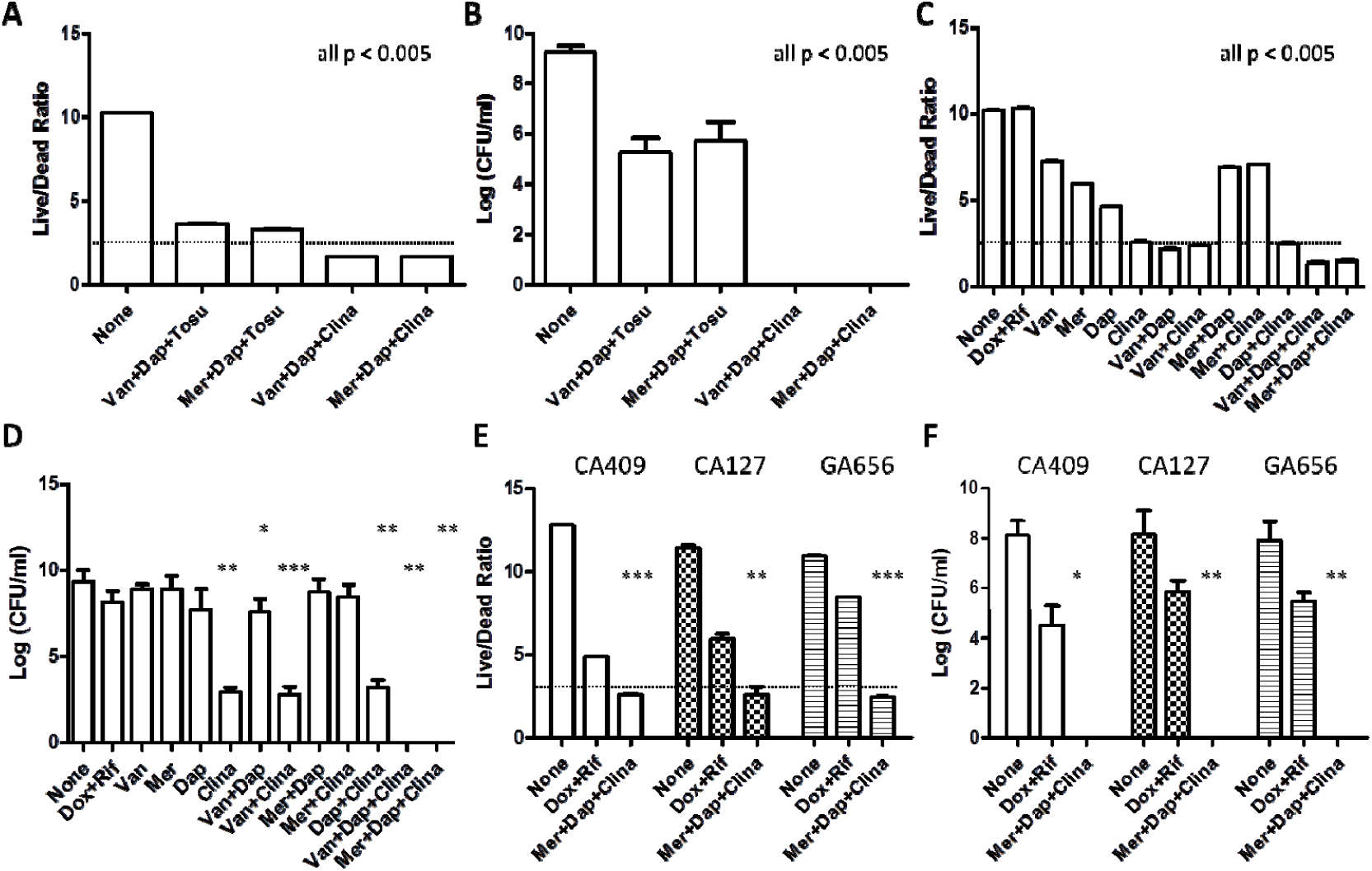
Identification of drug combinations that kill MRSA biofilms. Various drug combinations consisting of drugs (at 50 μM) highly active against growing phases and persister cells were tested and evaluated for their anti-biofilm activity by SYBR Green I/PI viability staining (A) and CFU enumeration (B). Drug combinations with sterilizing activity against USA300 biofilms were tested at clinically achievable concentrations (Cmax). (E,F) Validation of meropenem + daptomycin + clinafloxacin in killing biofilms of various clinical isolates of MRSA (C,D). Vancomycin, Van; Meropenem, Mer; Daptomycin, Dap; Tosufloxacin, Tosu; Clinafloxacin, Clina; Doxycycline, Dox; Rifampin, Rif. Student’s *t*-test, \**p*□<□0.05, \*\**p*□<□0.005, \*\*\**p*□<□0.0005.

We then tested the potential of the drug combination of meropenem + daptomycin + clinafloxacin to eradicate biofilm bacteria from different MRSA *S. aureus* strains, including other CA-MRSA clinical isolates CA-409, CA-127, and hospital-acquired MRSA strain GA-656. Complete eradication (0 CFU/ml) and undetectable levels of live cells (under the limit of detection) were found for all of the MRSA strains tested after 4 days of treating biofilms with our combination meropenem + daptomycin + clinafloxacin (Fig. 2E-F).

### Unique anti-persister activity of clinafloxacin that could not be replaced by other fluoroquinolone drugs

Clinafloxacin is a member of the fluoroquinolone class of antibiotics which inhibits DNA replication by binding to DNA gyrase. As our results suggest (Fig. 1), clinafloxacin is a powerful anti-persister drug. We then wanted to rank the anti-biofilm activity of different fluoroquinolones to determine whether the robust anti-biofilm activity of clinafloxacin used in combination is unique to the drug itself or can be replaced by other members of fluoroquinolone antibiotics. To do so, we used the *S. aureus* Newman strain due to its susceptibility to many fluoroquinolones as we wanted to eliminate any confounding factors due to inherent drug resistance. While other fluoroquinolones such as ciprofloxacin, levofloxacin, and moxifloxacin had certain anti-persister or anti-biofilm activity when used in combination with meropenem and daptomycin after 4-days of treatment, the drug combination with clinafloxacin was indeed the most active and was the only combination that achieved complete sterilization. By contrast, biofilms treated with combinations consisting of other quinolones still harbored 10^4^-10^8^ CFU/ml. When used in combination, the activity of the quinolones from strongest to weakest as ranked by both viability assessment and viable cell counts is as follows: clinafloxacin, ciprofloxacin, moxifloxacin, and levofloxacin (Table 2). Hence, clinafloxacin has unique potent activity against persisters compared to other fluoroquinolone counterparts.

**Table 2.** Ranking of Fluoroquinolones for Their Activity in Killing Biofilm Bacteria.

| **Treatment (Cmax)** | **Log**<br>**(CFU/ml)** | **Live/Dead Ratio** | **Ranking of**<br>**combination**<br>**(1= best,**<br>**4= worst)** |
| --- | --- | --- | --- |
| **None** | 9.23 ± 0.4 | 11.95 ± 0.1 | n/a |
| **Meropenem** | 7.34 ± 0.7 | 8.11 ± 0.04 | n/a |
| **Daptomycin** | 9.34 ± 0.3 | 7.92 ± 0.2 | n/a |
| **Meropenem+Daptomycin** | 6.77 ± 0.6 | 4.52 ± 0.03 | n/a |
| **Mer+Dap+Clina** | 0 | 1.03 ± 0.07 T | 1 |
| **Mer+Dap+Cipro** | 3.87 ± 0.3 | 1.34 ± 0.08 T | 2 |
| **Mer+Dap+Moxi** | 4.99 ± 0.6 | 1.24 ± 0.06 T | 3 |
| **Mer+Dap+Levo** | 8.91 ± 0.5 | 7.19 ± 0.1 T | 4 |

T = Limit of detection has been reached for the SYBR Green I/PI viability assay
Meropenem, Mer; Daptomycin, Dap; Clinafloxacin, Clina; Ciprofloxacin, Cipro; Moxifloxacin, Moxi; Levofloxacin, Levo
n/a= none available

### Anti-persister activity of oritavancin and dalbavancin

Thus far, our data suggest that inclusion of a drug with great anti-persister activity can be beneficial in killing biofilm bacteria (Fig. 1, Fig. 2, and Table 2). To identify other potential anti-persister candidates, we turned to the new generation of lipoglycopeptides such as oritavancin and dalbavancin. These drugs have multiple mechanisms of action: inhibition of transglycosylation, transpeptidation, and cell membrane disruption, a property of persister drugs^19^.We first tested the activity of oritavancin and dalbavancin in killing *S. aureus* persisters in comparison with its parent counterpart vancomycin, and the results revealed oritavancin was the best in killing persisters among the three drugs (Fig. 3A). After 6-day drug exposure, oritavancin killed 10^6^ CFU/ml of persisters as compared with dalbavancin or vancomycin which killed only about 10^2^ CFU/ml.

**Figure 3.**
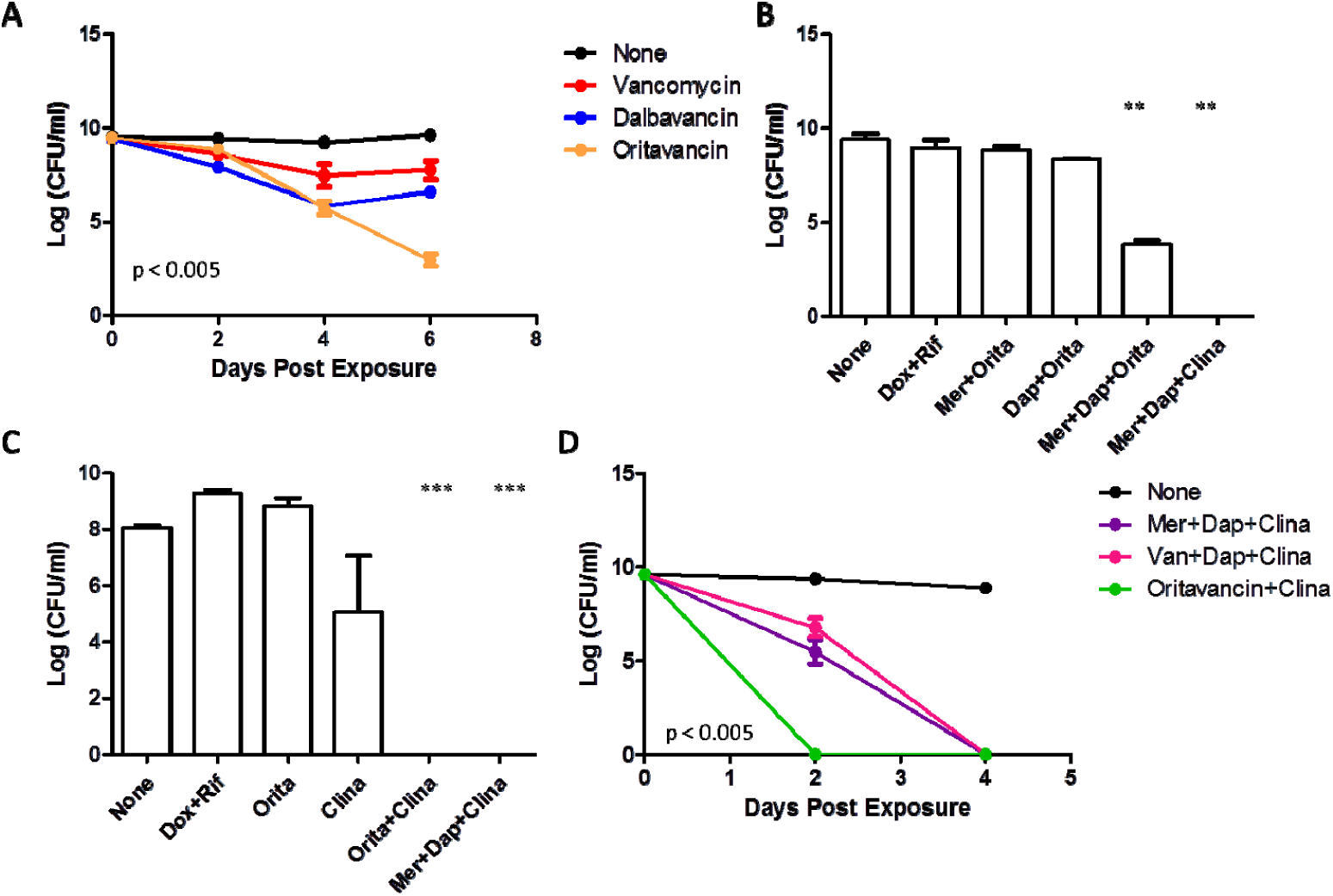
Evaluation of oritavancin in killing biofilms as a single drug or in combination. (A) Comparing novel lipoglycopeptides oritavancin and dalbavancin to vancomycin in their activity to kill persisters (at 50 μM). Evaluating oritavancin in (B) killing persisters in combination with meropenem+daptomycin and the agent (C) killing growing phase bacteria in combination with clinafloxacin (at Cmax concentrations). (D) Time-kill curve of biofilms comparing the top drug combinations candidates (at Cmax concentrations). Meropenem, Mer; Daptomycin, Dap; Oritavancin, Orita; Clinafloxacin, Clina; Doxycycline, Dox; Rifampin, Rif. Two-way ANOVA, and Student’s *t-*test, *p□<□0.05, **p□<□0.005, ***p□<□0.0005.

Since oritavancin showed strong anti-persister activity, we next evaluated oritavancin’s activity in drug combinations. After replacing clinafloxacin with oritavancin, we observed that the combination of meropenem + daptomycin + oritavancin exhibited partial activity against biofilms, a decrease of 10^5^ CFU/ml, which is much better than the activity achieved by treatment with single drugs or two-drug combinations, but still inferior to the clinafloxacin combination (Fig. 3B).

Due to oritavancin’s strong activity against growing phase *S. aureus* (MIC of 0.03 mg/L) ^30^ and its dual mechanism of action that mimics cell wall + cell membrane inhibitors in our drug combination, we tested oritavancin in place of meropenem and daptomycin. Surprisingly, the combination of oritavancin + clinafloxacin was also able to achieve complete eradication of biofilms suggesting that oritavancin can replace the component in our drug combination that targets actively growing bacteria (Fig. 3C). It is also important to note that single drug of oritavancin cannot kill biofilms (no change in CFU after 4 day treatment) which further validates the importance of drug combinations in biofilm bacteria.

To compare the activity of the three combinations tested thus far with clinafloxacin, we performed a time-course kill experiment which revealed that oritavancin + clinafloxacin can kill all biofilms by 2-day treatment whereas it took 4 days for meropenem/vancomycin + daptomycin + clinafloxacin to eradicate the biofilm bacteria (Fig. 3D). Overall, our data suggest that inclusion of an anti-persister drug in a drug combination to treat biofilms is paramount and these combinations possess better activity than current clinically used regimens based on our in vitro studies.

### *The drug combination meropenem + daptomycin +clinafloxacin eradicated biofilm infections in the mouse skin persistent* infection model

Given the robust activity of our drug combinations in eradicating biofilms *in vitro*, we were interested to know if our combination can also eradicate persistent infections *in-vivo*. To evaluate the efficacy of our drug combinations in treating the persistent skin infection, we chose to infect mice with biofilm bacteria from *S. aureus* strain USA300, a clinical strain most representative to causing persistent infections in a host. We allowed the infection to develop for 7 days, followed by 7 day treatment with different regimens (Fig. 4A). Previously, we have shown that mice infected with biofilm bacteria developed more chronic skin lesions ^31^. Administration of the combination of doxycycline + rifampin (a control group as a clinically used treatment) or drug combination vancomycin + daptomycin + clinafloxacin decreased the bacteria load (about 1-log of bacteria) but did not clear the infection (Fig. 4B). Other treatments which supposedly eradicate chronic *S. aureus* infections such as ADEP4+rifampin ^32^ or frustose+gentamicin ^33^ did not show sterilizing activity in our biofilm infection model and instead, had increased lesion size and inflammation (Fig. 4C). Remarkably, our combination of meropenem + daptomycin + clinafloxacin cleared the infection completely, decreased the size of lesions, and reduced histopathology scores, and healed the lesions completely (Fig. 4D-G).

**Figure 4.**
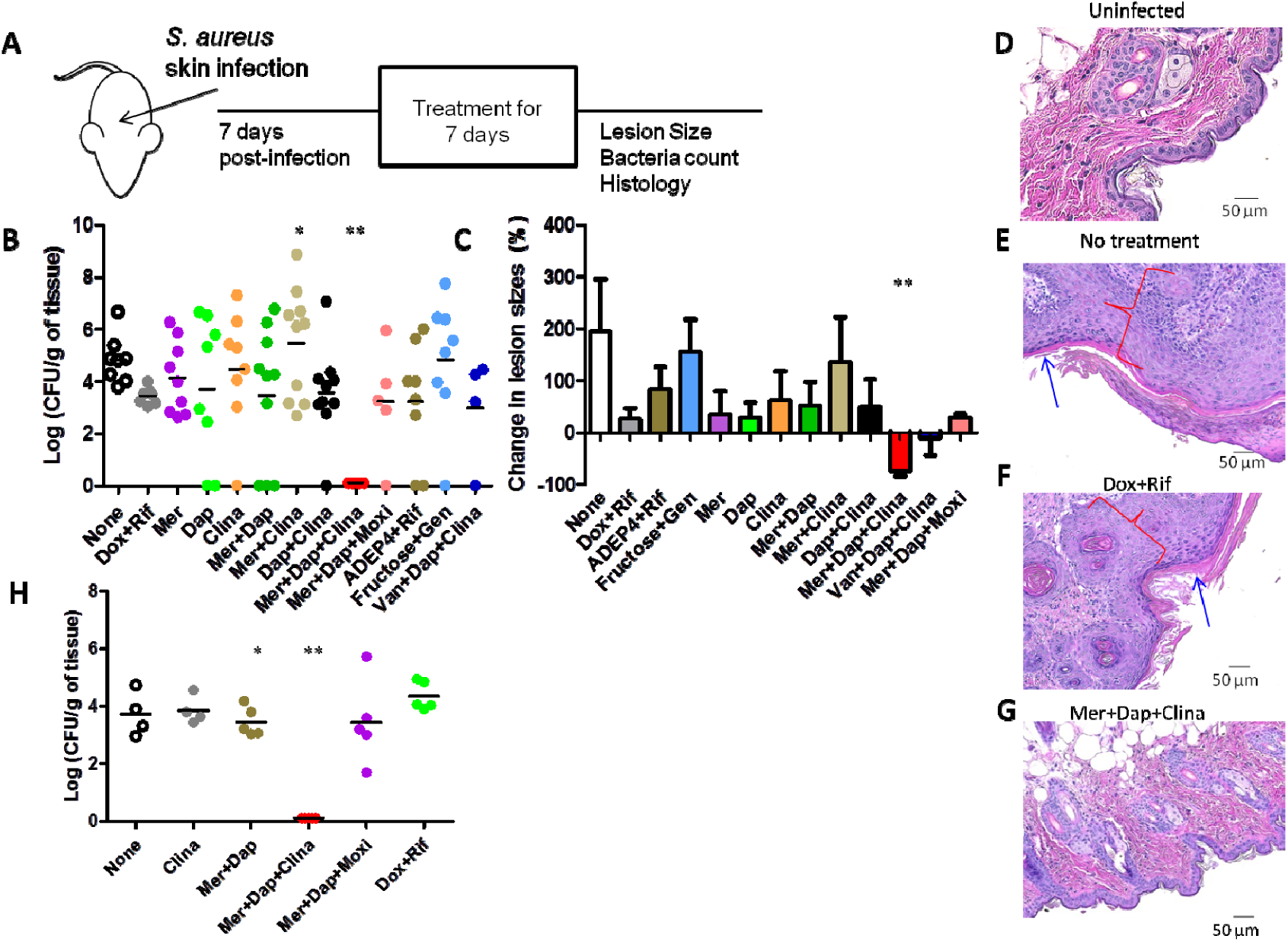
Validation of drug combinations in chronic skin infection model. (A) Study design of mouse treatment studies. (B) Bacterial load in the skin lesions. (C) Changes in lesion sizes and histology of skin tissues of mice infected with USA300 and treated 7-days with drug combinations and respective controls were measured. Histopathology of uninfected mice (D), infected mice receiving 7-days of no treatment (E), or treated with doxycycline + rifampin (F), or treated with meropenem + daptomycin + clinafloxacin (G) was analyzed. (H) Bacterial loads in the skin tissues of mice infected with MSSA Newman strain and treated with 7-days with drug combinations or control treatments were enumerated. Images were taken at 200X magnification. Meropenem, Mer; Daptomycin, Dap; Tosufloxacin, Tosu; Clinafloxacin, Clina; Doxycycline, Dox; Rifampin, Rif, Gentamicin, Gen. Blue arrows indicate crust formation, red brackets indicate hyperplasia and cellular infiltration; Student’s *t*-test, \**p*□<□0.05, \*\**p*□<□0.005, \*\*\**p*□<□0.0005.

Because our in-vivo experiments were done using an MRSA strain USA300, we wanted to infect mice with a methicillin susceptible *S. aureus* Newman strain (MSSA) to ensure that our drug combination is effective for other *S. aureus* strains. In addition, we wanted to further confirm that clinafloxacin’s activity, compared to other quinolones, is superior despite the difference in background sensitivity of the bacterial strain. Despite moxifloxacin and clinafloxacin having the same MIC for the Newman strain, the combination of meropenem + daptomycin + moxifloxacin was not effective in clearing the biofilm infection whereas the combination of meropenem + daptomycin + clinafloxacin indeed cleared the infection completely (Fig. 4H). This indicates clinafloxacin combination works for both MRSA strain and MSSA strain and its unique sterilizing activity cannot be replaced by moxifloxacin.

### Eradication of biofilm bacteria correlates with resolution of tissue inflammation and immunopathology

Skin infections caused by *S. aureus* are cleared by neutrophil recruitment driven by IL-17 production ^34^. To evaluate any potential immunopathology consequences of our treatment, we measured the levels of IL-17 and proinflammatory cytokine IL-1 at the infection site and performed histology. Skin tissues of mice treated with our drug combination of meropenem + daptomycin + clinafloxacin produced the lowest amount of IL-17 (Fig. 5A) and IL-1 (Fig. 5B) and showed the least amount of gross inflammation (Fig. 5C) compared to other control treatments. These data support our hypothesis that a drug combination targeting both growing (e.g. meropenem, daptomycin) and persister cells (e.g. clinafloxacin) is essential in clearing chronic infections *in-vivo* such as a persistent skin infection caused by *S. aureus* biofilm.

**Figure 5.**
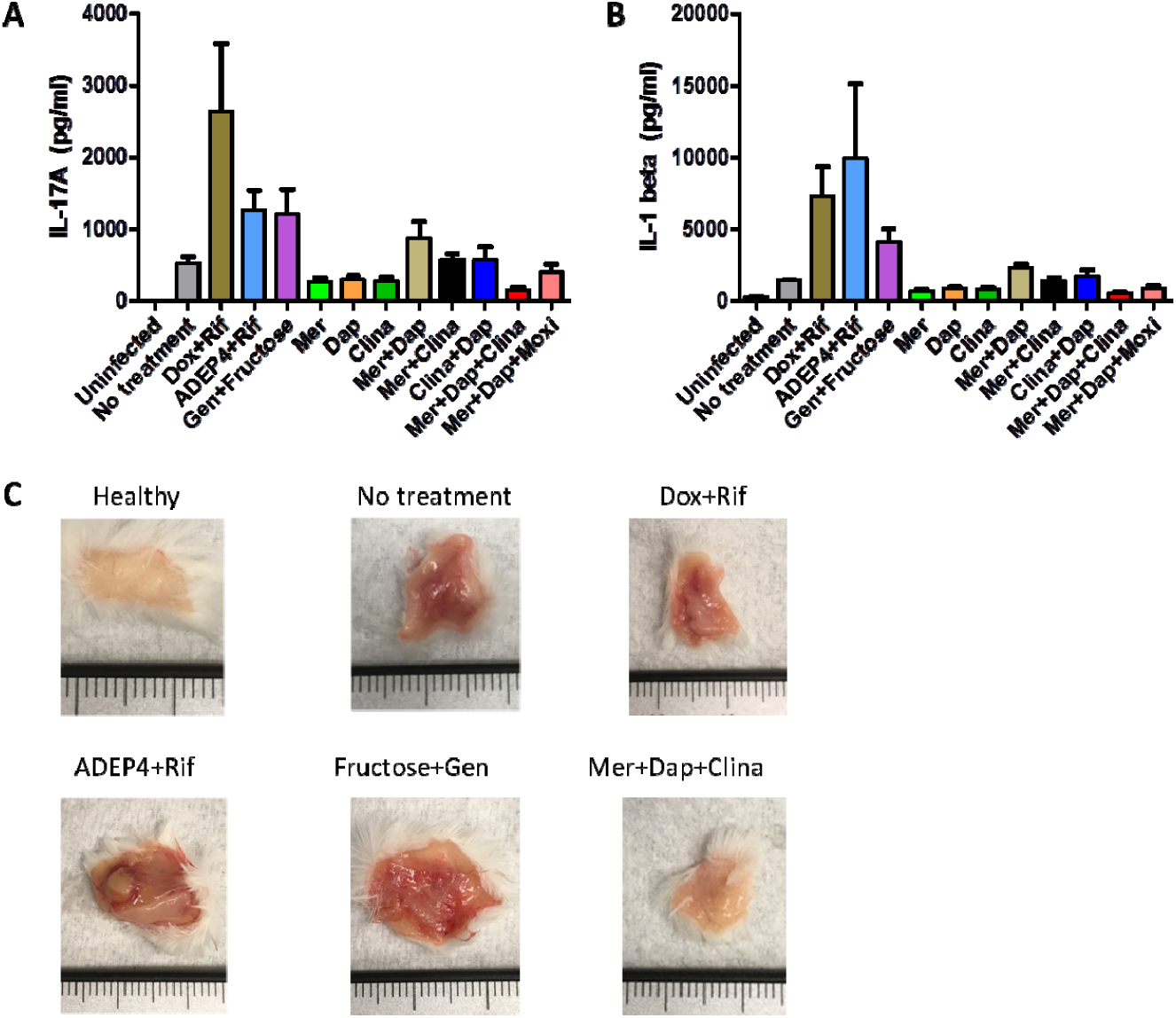
Meropenem + Daptomycin + Clinafloxacin reduced immune response of infected skin tissues. (A) IL-17, (B) IL-1, and (C) Gross pathology of skin tissues of mice treated 7-days with drug combinations and respective controls (in mm). Meropenem, Mer; Daptomycin, Dap; Tosufloxacin, Tosu; Clinafloxacin, Clina; Doxycycline, Dox; Rifampin, Rif, Gentamicin, Gen

## Discussion

Numerous studies have documented how resilient biofilms and biofilm infections are to antibiotic treatments ^35-37^Since persister cells that are embedded in the biofilm are mostly responsible for recalcitrance of biofilms to antibiotic treatments, many attempts have been made to identify novel treatments and synthetic compounds that kill bacterial persisters ^38,39^. Some approaches include resuscitating or altering the metabolic status of persisters ^40,41^or enhancing the activity of aminoglycoside antibiotic with sugars ^33^, or activating protease by ADEP4 plus rifampin^32^. Although these new therapeutic approaches showed promising results *in vitro* and in some cases, *in-vivo*, ^32,38,42^ not all treatments achieved sterilizing activity and their utility in more persistent biofilm infections remains to be confirmed. The animal models used either rely on immunosuppressant agents or a short term infection that do not reflect true persistent infections clinically. Here, we established a more relevant persistent infection mouse model with biofilm inocula that mimic the human infections without the use of immunosuppressant agents. Previous studies have mostly used log phase bacteria as inocula for infection in animal models, however, in this study, we showed that biofilm inocula produced a more severe lesion and more persistent infection than the log phase bacteria^31^. This biofilm-inocula model could serve as a useful model for evaluating treatment regimens against biofilm infections in vivo in general. Importantly, we were able to show that single drugs or even two drug combinations can this study, to identify more effective regimens to treat chronic *S. aureus* infections, we first identified several drug combinations that are more active in killing biofilms *in vitro* than currently recommended regimens (e.g. vancomycin alone, doxycycline + rifampin, and vancomycin + gentamicin + rifampin) used clinically. Then, we confirmed the potent activity of the combination meropenem + daptomycin + clinafloxacin in our newly established chronic, skin infection mouse model.

Previously, we identified both clinafloxacin and tosufloxacin as having robust activity against *S. aureus* persisters ^27^. However, in our drug combination studies, clinafloxacin used in combination displayed greater activity against biofilms than tosufloxacin. This could potentially be explained by the genetic background and inherent antibiotic resistance of the strain tested. Previously, tosufloxacin was identified to have great persister activity in the background of the Newman strain ^27^, a methicillin-sensitive *S. aureus* strain whereas our biofilm experiments conducted here used USA300, a MRSA strain. The ability between these strains to form biofilms may also be a contributing factor ^43^. Nonetheless, despite being unable to achieve complete eradication, it must be noted that inclusion of any anti-persister drug in combination (clinafloxacin, tosufloxacin, or oritavancin) with drugs that can kill growing bacteria can kill more bacteria in biofilms than currently approved regimens, confirming the importance of targeting the heterogeneous of bacterial populations in developing more effective treatments.

We showed that meropenem + daptomycin + clinafloxacin achieved sterilizing activity in a chronic, skin infection model in mice. The strong activity of meropenem or daptomycin against *S. aureus* growing bacteria is indisputable as most *S. aureus* strains have a relatively low MIC to these drugs ^44,45^. The inclusion of drugs with such strong activity against active bacteria allows for rapid killing of growing bacteria in the population. To kill non-growing biofilm persisters, clinafloxacin is crucial in the combination. As shown here, the MICs for clinafloxacin for both the MSSA Newman strain and MRSA USA300 strain were both under 0.25 µg/ml. Other studies have shown that MICs for streptococci are from 0.06-0.12 µg/ml and for enterococci is 0.5 µg/ml. Both oral and intravenous formulations have been developed ^46,47^. Although not commonly used, clinafloxacin administration drastically improved the condition of a cystic fibrosis patient who had a chronic *Burkholderia cenocepacia* infection and was not responding to different antibiotic treatment ^48^. A human trial with patients having native or prosthetic valve endocarditis also showed that clinafloxacin was an effective treatment ^49^. As a quinolone, clinafloxacin inhibits bacterial DNA gyrase and topoisomerase IV but not all quinolones have anti-persister activity (Table 1). Comparing the chemical structure of clinafloxacin to the other quinolones that have weak anti-persister activity (ciprofloxacin, tosufloxacin, moxifloxacin, and levofloxacin), a chloride group attached to the benzene ring appears to be unique to only clinafloxacin (Fig. 6). Further studies to explore the mechanism of clinafloxacin’s unique ability to kill persisters requires further investigation.

**Figure 6.**
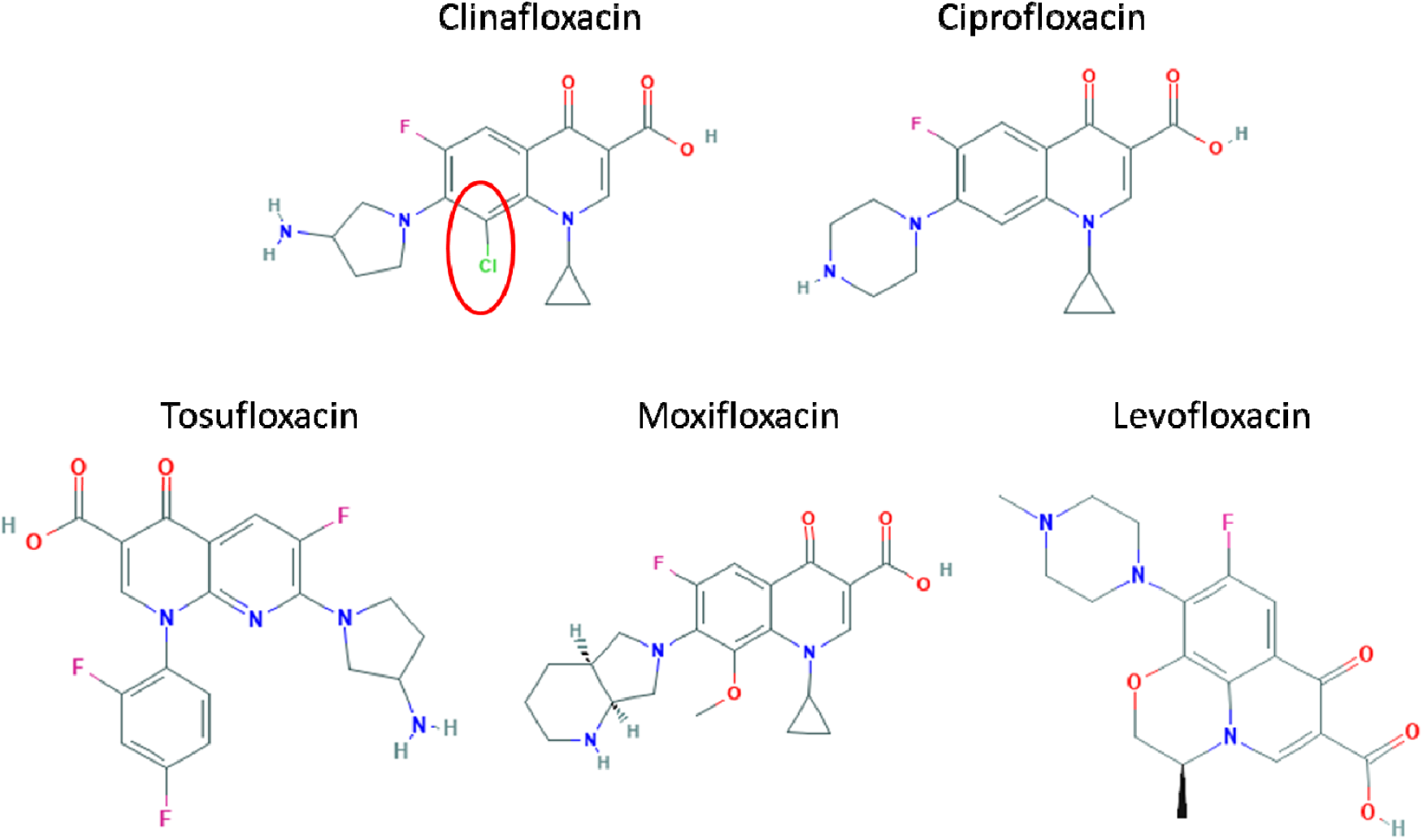
Chemical structures of the fluoroquinolones tested. The chloride group (circled) is unique to clinafloxacin.

While meropenem and daptomycin are our agents directed at killing growing bacteria, it is important to note that these drugs also have some activity against persisters. Meropenem used in combination with polymyxin B has been shown to eradicate persisters in *Acinetobacter baumannii* strains ^50^. Similarly, daptomycin has been shown to be active against *S. aureus* biofilms found on implants ^51^. Daptomycin in combination with doxycycline and cefoperazone or cefuroxime have been shown to kill biofilm-like microcolonies of *B. burgdorferi* ^*26,52*^. Its mechanism of action in disrupting membrane structure and rapid depolarization of the membrane may impact the viability of persisters and thus, play an important role in the combination ^53^.

It is important to note that the chronic infection status of our mice is a key component to our disease model. The phenotype of a more severe disease caused by persistent forms is an important observation ^31^ and our model could potentially better mimic chronic infections in humans. While Conlon et al. also used stationary phase inocula (10^6^) to infect their mice and caused a deep-seated infection, the infection was allowed to develop for only 24 hours before treatment which cannot be a persistent infection; and the mice were made neutropenic ^32^, a condition that may not apply to a majority of patients suffering from chronic *S. aureus* infections. Such differences in the animal models may explain why ADEP4 + rifampin which was claimed to have sterilizing activity by Conlon et al. failed to eradicate the persistent infection in our model established with biofilm inocula and infection allowed to develop for 1 week before treatment. The combination of an aminoglycoside + sugar was shown to be effective in an *E. coli* urinary tract infection model ^33^ but unfortunately this approach was not effective in our biofilm infection model either. Allison *et al.* showed that gentamicin + fructose reduced 1.5 fold of *S. aureus* biofilms *in vitro* after 4 hours of treament^33^ but was not tested in animals. In our study here, we showed that mice treated for 7 days with gentamicin + fructose still harbored 10^5^ CFU/ml in skin tissues and showed an increase in lesion size despite the treatment. In both these cases, the discrepancy could be due to differences in the disease model, as ours is a more persistent biofilm skin infection model established with biofilm inocula and would be expected to be more difficult to cure than the other studies that did not use biofilm inocula for the infection.

Our combination of meropenem + daptomycin + clinafloxacin showed sterilizing activity in mice after one week based on the concentrations of drugs and dosing regimens commonly found in literature (Table 2). Higher, yet safe, concentrations of drugs should be tested to see whether or not a shorter treatment period can achieve similar eradication, which would be beneficial for patients undergoing therapy. Further PK/PD studies of our drug combination are needed. Moreover, meropenem, daptomycin, and clinafloxacin are intravenous drugs and not convenient to administer. Future studies to develop oral regimens as effective as the identified combinations are needed for more convenient administration. Our *in vitro* data suggested that oritavancin used in combination with clinafloxacin had robust activity against biofilms, killing all the bacteria in the biofilms (10^7^ CFU) after a short treatment of 2 days. The administration of oritavancin is a single 1200-mg dose given in a slow, 3 hour infusion, which may also be of interest for patients due to the ease of administration and long half-life. Hence, preclinical studies in mice to test oritavancin’s activity in chronic infections need to be performed carefully.

Currently used regimens for treating persistent infections are lengthy and the inability to clear the bacteria in a timely fashion may also increase the chances of developing antibiotic resistance. A drug combination that has both activities against growing and persister cells as proposed in the Yin-Yang model^25^ have promising potential in developing a more effective therapy for treating chronic persistent infections. This study validates this Yin-Yang model^25^ and emphasizes the importance of persister drugs like clinafloxacin in eradicating a persistent infection. This treatment algorithm takes into account the heterogeneous population of bacterial cells that exists upon encountering stress. With this principle in mind, this study reports novel drug combinations that are effective in killing *S. aureus* biofilms and treating chronic infections. We established a chronic skin infection mouse model that more appropriately mimics human chronic disease. Then, we developed a triple drug combination of meropenem (or vancomycin) + daptomycin + clinafloxacin or two drug combination of oritavancin + clinafloxacin that can achieve sterilizing activity *in vitro*. We also show that administration of meropenem + daptomycin + clinafloxacin allowed the mice with chronic skin infections to completely clear the bacterial load, heal lesions completely, and show reduced pathology and inflammation. Our approach of combining drugs targeting both growing and non-growing bacteria with persister drugs to completely eradicate biofilm infections may have implications for developing better treatments against other persistent infections by other bacterial pathogens, fungi, and even cancer.

## Methods

### Culture media, antibiotics, and chemicals

*Staphylococcus aureus* strains Newman, USA300, CA-409, CA-127, and GA-656, were obtained from American Type Tissue Collections (Manassas, VA, USA), and cultivated in tryptic soy broth (TSB) and tryptic soy agar (TSA) (Becton Dickinson Franklin Lakes, NJ, USA) at 37°C. Vancomycin, gentamicin, rifampicin, levofloxacin, ciprofloxacin, moxifloxacin, and oritavancin were obtained from Sigma-Aldrich Co. (St. Louis, MO, USA). Daptomycin, meropenem, tosufloxacin, and clinafloxacin were obtained from AK Scientific, Inc. (Union City, CA, USA). Stock solutions were prepared in the laboratory, filter-sterilized and used at indicated concentrations.

### Microtiter plate biofilm assays

*S. aureus* strains grown overnight in TSB were diluted 1:100 in TSB. Then, 100-μl aliquots of each diluted culture were placed into a 96-well flat-bottom microtiter plate and statically incubated for 24 h at 37°C ^29^. Planktonic cells were removed and discarded from the microtiter plates. Drugs at the indicated concentrations were then added to the biofilms attached to the bottom of the microtiter plate at a total volume of 100 μl in TSB. To determine the cell and biofilm density, the supernatant was removed from the well and the biofilms were washed twice with PBS (1X). To enumerate bacterial cell counts, the biofilms in the wells were resuspended in TSB and scraped with a pipette tip before serial dilution and plating. To assess cell viability using the ratio of green:red fluorescence to determine the ratio of live:dead cells, respectively, the biofilms were stained with SYBR Green I/Propidium Iodide dyes as described ^28,54^. Briefly, SYBR Green I (10,000× stock) was mixed with PI (20 mM) in distilled H_2_O at a ratio of 1:3, respectively. The SYBR Green I/PI staining mix was added to each sample at a ratio of 1:10 (10 µl of dye for 100 µl of sample). Upon incubation at room temperature in the dark for 20 min, the green and red fluorescence intensity was detected using a Synergy H1 microplate reader by BioTek Instruments (Winooski, VT, USA) at excitation wavelength of 485 nm and 538 nm and 612 nm for green and red emission, respectively. To visualize biofilm biomass, biofilms were stained with crystal violet (0.1%) for 15 minutes at room temperature. Excess dyes were washed with water and the biofilms were left to air dry. Images were recorded using Keyence BZ-X710 microscope and were processed using BZ-X Analyzer software provided by Keyence (Osaka, Japan).

### Mouse skin infection model

Female Swiss-Webster mice of 6 weeks of age were obtained from Charles River. They were housed 3 to 5 per cage under BSL-2 housing conditions. All animal procedures were approved by the Johns Hopkins University Animal Care and Use Committee. *S. aureus* strain USA300 and strain Newman were used in the mouse infection experiments. Mice were anesthetized and then shaved to remove a patch of fur of approximately 3 cm by 2 cm. Bacteria of indicated inoculum size and age were subcutaneously injected into the mice. For log phase inoculum, bacteria grown overnight were diluted 1:100 in TSB and grown for 2 hrs at 37°C with shaking at 220 RPM. For stationary phase inoculum, overnight cultures of bacteria grown at 37°C were used. For preparation of biofilm inoculum, biofilms were first grown in microtiter plates as described previously, and then resuspended and scraped up with a pipette tip. Quantification of all inoculum was performed by serial dilution and plating. Treatment was started after 1 week infection with different drugs and drug combinations. For details on drugs, drug dosage and route of administration, please refer to Table 1. Skin lesion sizes were measured at the indicated time points using a caliper. Mice were euthanized after 1 week post-treatment and skin tissues were removed, homogenized, and serially diluted for bacterial counting on TSA plates.

### Histopathology

Skin tissues were dissected, laid flat, and fixed for 24 hrs with neutral buffered formalin. Tissues were embedded in paraffin, cut into 5-μm sections, and mounted on glass slides. Tissue sections were stained with hematoxylin and eosin for histopathological scoring. Tissue sections were evaluated for lesion crust formation, ulcer formation, hyperplasia, inflammation, gross size, and bacterial count and were assigned a score on a 0–3 scale (0 = none, 1 = mild, 2 = moderate, and 3 = severe). The cumulative pathology score represented the sum of each individual pathology parameter. Scoring was performed by an observer in consultation with a veterinary pathologist. Representative images were taken using a Keyence BZ-X710 Microscope.

### Statistical analyses

Statistical analyses were performed using two-tailed Student’s *t*-test and two-way ANOVAs where appropriate. Mean differences were considered statistically significant if *p* value was <0.05. All experiments were performed in triplicates. Analyses were performed using GraphPad Prism and Microsoft Office Excel.

